# Overlapping transcriptional expression response of wheat zinc-induced facilitator-like transporters emphasize important role during Fe and Zn stress

**DOI:** 10.1101/581405

**Authors:** Shivani Sharma, Gazaldeep Kaur, Anil Kumar, Jaspreet Kaur, Ajay Kumar Pandey

**Affiliations:** National Agri-Food Biotechnology Institute (Department of Biotechnology), Sector 81, Knowledge City, Mohali, Punjab, India-140306; University Institute of Engineering and Technology, Sector 25, Panjab University, Chandigarh, Punjab, India-160015

**Keywords:** iron starvation, micronutrient uptake, *Triticum aestivum* L., zinc transport, biofortification

## Abstract

**Background:** Hexaploid wheat is an important cereal crop that has been targeted to enhance grain micronutrient content including zinc and iron. In this direction, modulating the expression of plant transporters involved in Fe and Zn homeostasis could be one of the promising approaches. Therefore, the present work was undertaken to identify bread wheat <u>Z</u>inc–Induced <u>F</u>acilitator-<u>L</u>ike (ZIFL) family of transporters and study their transcriptional expression response during micronutrient fluctuations and exposure to multiple heavy metals.

**Results:** The genome-wide analyses resulted in identification of thirty-five putative *TaZIFL* genes, which were distributed only on Chromosome 3, 4 and 5. Wheat ZIFL proteins subjected to the phylogenetic analysis showed the uniform distribution along with rice, *Arabidopsis* and maize. *In-silico* analysis of the promoters of the wheat ZIFL genes suggested the presence of multiple metal binding sites including those which are involved in Fe homeostasis. QRT-PCR analysis of wheat *ZIFL* genes suggested the differential regulation of the transcripts in roots and shoots under surplus Zn and also during Fe starvation. Specifically, in roots, *TaZIFL2.3, TaZIFL4.1, TaZIFL4.2, TaZIFL5, TaZIFL6.1* and *TaZIFL6.2* were significantly up-regulated by both Zn and Fe. This suggested that ZIFL could possibly be regulated by both the nutrient stress in a tissue specific manner. Interestingly, upon exposure to heavy metals, *TaZIFL4.2* and *TaZIFL7.1* showed significant up-regulation, whereas *TaZIFL5* and *TaZIFL6.2* remained almost unaffected.

**Conclusion:** This is the first report with detailed analysis of wheat ZIFL genes. Our study also identifies closest ortholog for transporter of mugineic acid, a chelator required for Fe uptake. Comprehensive transcript expression pattern during development of wheat seedlings and against various abiotic/biotic stresses resulted in tissue specific responses. Overall, this work addresses the role of wheat *ZIFL* during the interplay between micronutrient and heavy metal stress in a tissue specific manner.

## Background

Crop plants are an important target for enhancing micronutrient content, including iron (Fe) and zinc (Zn) in the developing grains. From human nutritional point of view, these micronutrients play an important role in growth and development, cognitive and immune impairment, and in gene regulation (Nyaradi et al., 2013; Gibson et al., 2018; Wishart et al., 2017). Therefore, researches worldwide are identifying diverse approaches to generate micronutrient rich food crops. Apart from a nutritional point of view, Fe and Zn are also essential minerals for plant development and various biochemical functions (Rout et al., 2015; Hafeez et al., 2013). Multiple metal specific transporters and regulators show co-expression and could share common signaling features including the response towards Zn, Fe or other metals. Several specific transporters are involved in the uptake and translocation of Fe and Zn, inside the plant has been described previously (Kawachi et al., 2018; Stein et al., 2009; Klein et al., 2009; Eide et al., 2014; Lee et al.,2009; Morrisey et al., 2009; Morel et al., 2008). Efficient micronutrient uptake by plants is a concerted effort of major genes belonging to the different families of transporters that include, but are not limited to zinc–regulated transporter, iron–regulated transporter family, the natural resistance associated macrophage protein family, yellow-stripe 1-like (YSL) subfamily of the oligopeptide transporter superfamily, Ca^2+^-sensitive cross complementer 1 (CCC1) family (Yoneyama et al., 2015; Ishimaru et al.,2011; Romheld et al., 2004; Sinclair et al.,2012; Kumar et al., 2018).

Many evidences are gathering that suggest an important role of major facilitator superfamily (MFS) clan of transporters, yet the identification of specific candidate genes has been always a bottleneck. Earlier, one such important group of genes referred as Zinc-induced facilitator-1 like gene (ZIFL), was identified and its role was assessed in different stresses including Zn homeostasis (Haydon et al., 2007). Role of ZIFL was not addressed in crop plants until the reports describing the inventory from model plant *Arabidopsis thaliana* and crop like *Oryza sativa* came forth (Haydon et al.,2007; Ricachenevsky et al., 2011). Since, the first identification of the three *ZIFL* in *Arabidopsis* referred to as *AtZIF1* (AT5G13740), *AtZIFL1* (AT5G13750) and *AtZIFL2* (AT3G43790) evidences are accumulating for their role specifically in Zn homeostasis (Haydon et al., 2007). Subsequently, multiple *ZIFL*s from monocot such as rice were identified and the presence of high numbers of these genes was correlated with the genome duplication events. Further, it was speculated that plant *ZIFL*s might perform redundant function that is imperative by their overlapping expression response for Fe and Zn (Haydon et al., 2012). In addition to that rice ZIFL family of genes were also characterized to be transporter of mugineic acid, a phytosiderophores involved in strategy-II mode of Fe uptake via roots (Nozoye et al.,2011; Nozoye et al., 2015). Recently, ZIFL1.1 in *Arabidopsis* was shown to impact the cellular auxin efflux at the root tip and contribute for drought tolerance (Remy et al., 2013). Furthermore, the role of ZIFLs was expanded for their involvement in potassium homeostasis and their ability to transport transition metal like cesium. (Remy et al., 2013). Subsequently, the maize (*Zea mays*) Zm-mfs1 was also identified and characterized. (Simmons et al., 2003). Such studies provide a valuable clue to the importance of ZIFL that is not just limited to transport of important micronutrients.

Hexaploid wheat (*Triticum aestivum* L.) is an important crop for the developing countries where, the suboptimal levels of grain Zn and Fe have been reported. Initiatives to enhance multiple micronutrients including Zn and Fe are being undertaken by either gain of function approach or RNAi mediated gene silencing (Connorton et al., 2017; Singh et al., 2017; Aggarwal et al., 2012). Earlier, using wheat grain transcriptome data it was confirmed that a higher proportion of transcripts were present in abundant amount that are known to be involved in transport activity (GO:0005215) (Gillies et al., 2012). Therefore, such studies provided the framework to investigate new resources and genes that could be of immense value to address the uptake and remobilization of micronutrients in cereal grains such as wheat. To provide impetus in this direction, the inventory of wheat ZIFL was built and their detailed expression characterization was performed.

In the current work, thirty-five wheat (*Triticum aestivum* L.) putative ZIFL proteins were identified that show their restricted distribution only on three chromosomes viz. 3, 4 and 5. Phylogenetic analysis revealed a uniform distribution of wheat ZIFL sequences in multiple clades along with rice, *Arabidopsis* and *Z. mays*. Detailed characterization of the *ZIFL* genes for their motif composition, promoter sequences and their expression under Fe limiting and Zn surplus condition and other heavy metals was also performed. Our data indicate that the wheat ZIFL show overlapping expression response during Fe starvation and Zn excess condition. Few of the wheat ZIFL gene expression remained unaffected by the presence of heavy metals. Overall, characterization of crop ZIFL transporters could result in identifying specific candidate/s that could be used further to modulate specific Fe-Zn uptake in crop plants such as wheat.

## Results

### Inventory of wheat ZIFL and their phylogeny analysis

In order to identify wheat *ZIFL* genes and to gain insight for possible evolutionary relationship, two complementary approaches were used. This includes, first performing genome-wide sequences of MFS_1 family using Pfam (PF07690) search, followed by homology-based analysis with previously reported *ZIFL* genes in different plant species Ensembl database. These approaches resulted in the identification of one hundred seventy-nine sequences and to further validate their identity sequences were checked and searched for a MFS_1 domain through Pfam and conserved domain databases (CDD-NCBI) (Table S2). These sequences were then used to build phylogenetic tree with previously known ZIFL proteins sequences from different plants (Table S1 and Figure S1). The arrangement of tree suggested a distinct clade for the ZIFL clustered when compared to the remaining MSF_1 proteins. This indicates that ZIFL is a distinct group of MFS transporters that are tightly clustered (Figure S1). Further, this distribution was validated through two signature sequences that are specific to ZIFLs (i) W-G-x(3)-D-[RK]-x-G-R-[RK] (except in TaZIFL2.5-5D) (ii) S-x(8)-[GA]-x(3)-G-P-x(2)-G-G with an exception of A instead of G at 10^th^ position of (ii) signature in TaZIFL2. Furthermore, sequences similar to ZIFL specific cysteine (Cys) and histidine (His) signatures were also used for identifying TaZIFLs. (iii) C-[PS]-G-C, absent in 5 TaZIFLs (TaZIFL2. TaZIFL 5-5D, TaZIFL 3-4B, TaZIFL 5-5D, TaZIFL 7.1-4B, TaZIFL 7.2-4B) probably due to missing sequence information and (iv) [PQ]-E-[TS]-[LI]-H-x-[HKLRD] (an insertion of ETLYCRHEHRYSIFISLD sequence within the motif was found in TaZIFL7.2-4A) (Ricachenevsky et al., 2011). The presence of one or another ZIFL signature motif further validated the distribution of genes. These signatures guided identification of specific wheat ZIFLs from the rest of the MFS_1 members. Such analysis resulted in confirmation of a total of thirty-five wheat ZIFL sequences, including individual *TaZIFL* genes and their respective homoeologos from different wheat sub-genomes (Table S3). To check the distribution along with other plant species, ZIFL protein sequences from *O. sativa, Z. mays* and *Arabidopsis* were used to build a rooted phylogenetic tree through the NJ method (Figure 1). Because of the genome duplication events in wheat, the genes are likely to show multiple alleles of a single gene. Hence the resulted thirty-five putative wheat *ZIFLs* represent 15 genes after distribution of their respective homoeologous (Figure 1). To provide the uniform nomenclature, *TaZIFL* genes were named according to their respective closest known orthologs from rice. Among the wheat ZIFL proteins TaZIFL4.1 showed highest homology with TaZIFL4.2 of 95.2 percentage identity. When a cross species comparison was done, the maximum identity of 87 percent was shown byTaZIFL2.2-3D and AtZIFL2. With rice, the highest percentage identity of 87 was observed for wheat ZIFL2.2-3A and OsZIFL2. The divergence was observed among TaZIFL3-4B and OsZIFL13 with percentage identity of 50.

**Figure 1.**
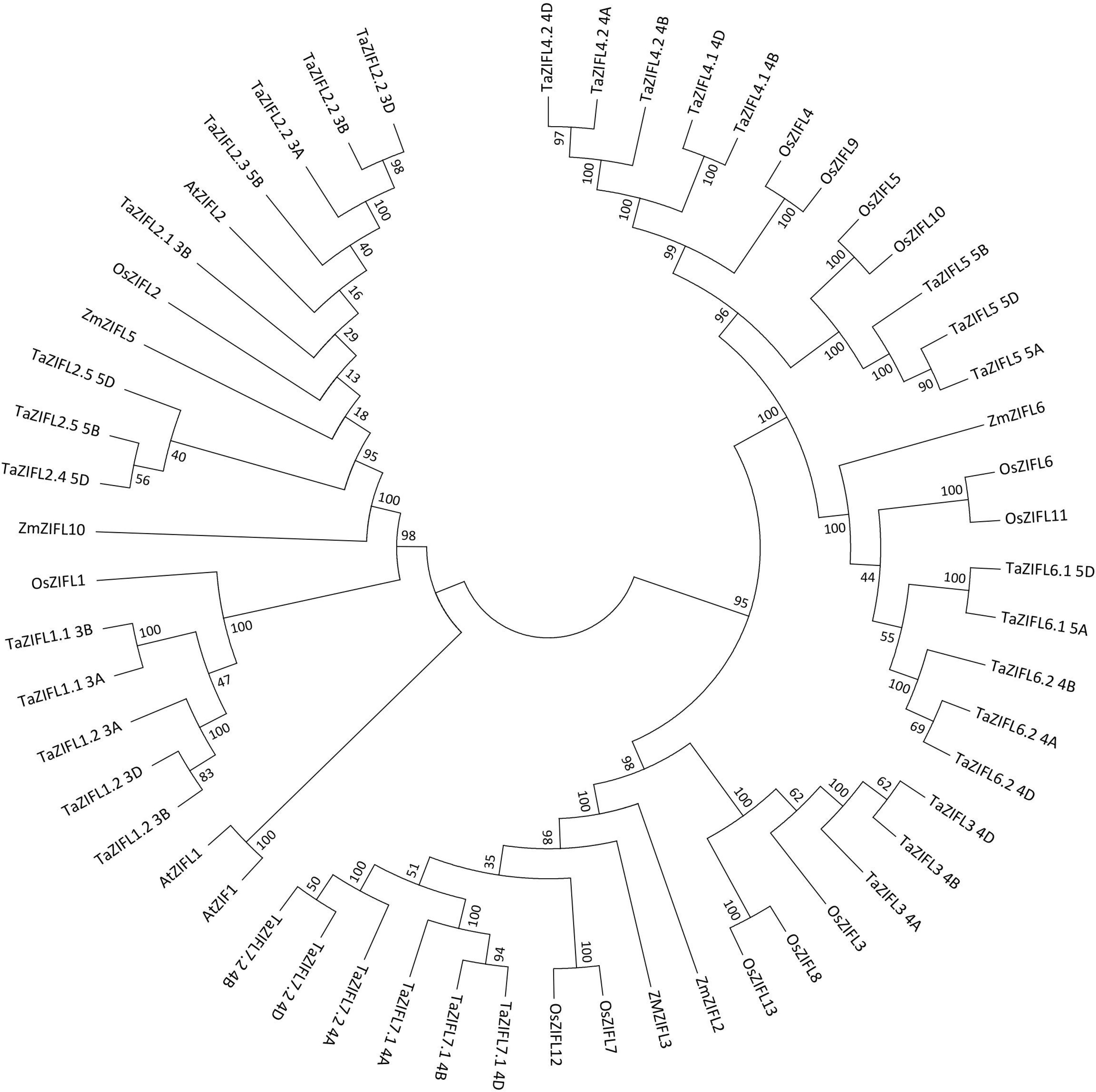
Phylogenetic analysis of wheat ZIFL protein sequences. The analysis was performed by using thirty-five ZIFL protein sequences from wheat, thirteen from *Oryza sativa*, seven from *Zea mays* and three from *Arabidopsis*. Rooted phylogenetic tree was constructed by using Neighbour-joining method using MEGA7 software with 1000 bootstrap replicates.

### Molecular structure and genome organization

The predicted protein length of the identified wheat ZIFL sequence ranged from 300 to 562 amino acids (Table S2). In general, most of the wheat ZIFL showed 10-12 predicted trans-membrane (TM) domains as reported in rice [19]. Specifically, 16 wheat ZIFL proteins were predicted to have 12 TM domains, 15 proteins have 10-11 predicted TM domains, 4 proteins were found to have 8-9 TM domains and only one wheat ZIFL has 4 TM domains (Table S2). Further, the genomic organization, analysis revealed the presence of genes on all three A, B and D sub-genomes. Maximum number of genes were found to be present on B and D sub-genome with 13 and 12 genes respectively (Figure 2a). TaZIFL1.2, TaZIFL2.2, TaZIFL3.4, TaZIFL5, TaZIFL6.2, TaZIFL7.1, TaZIFL7.2 are present in all three genomes, while TaZIFL2.3 and TaZIFL2.4 are present on only one genome 5B and 5D respectively (Table S2). The chromosomal distribution mapping revealed *TaZIFLs* to be present only on chromosome 3, 4 and 5 with maximum of 17 sequences on chromosome 4 (Figure 2b and c). Next, the genomic structure was analyzed and regions corresponding to intron-exons were marked (Figure 3). *TaZIFL* clustered into the same group and shared almost similar distribution pattern for the number of exon/intron. The intron-exon number varies from 14-17 in the respective *TaZIFL* genomic sequences (Figure 3, Table S2).

**Figure 2.**
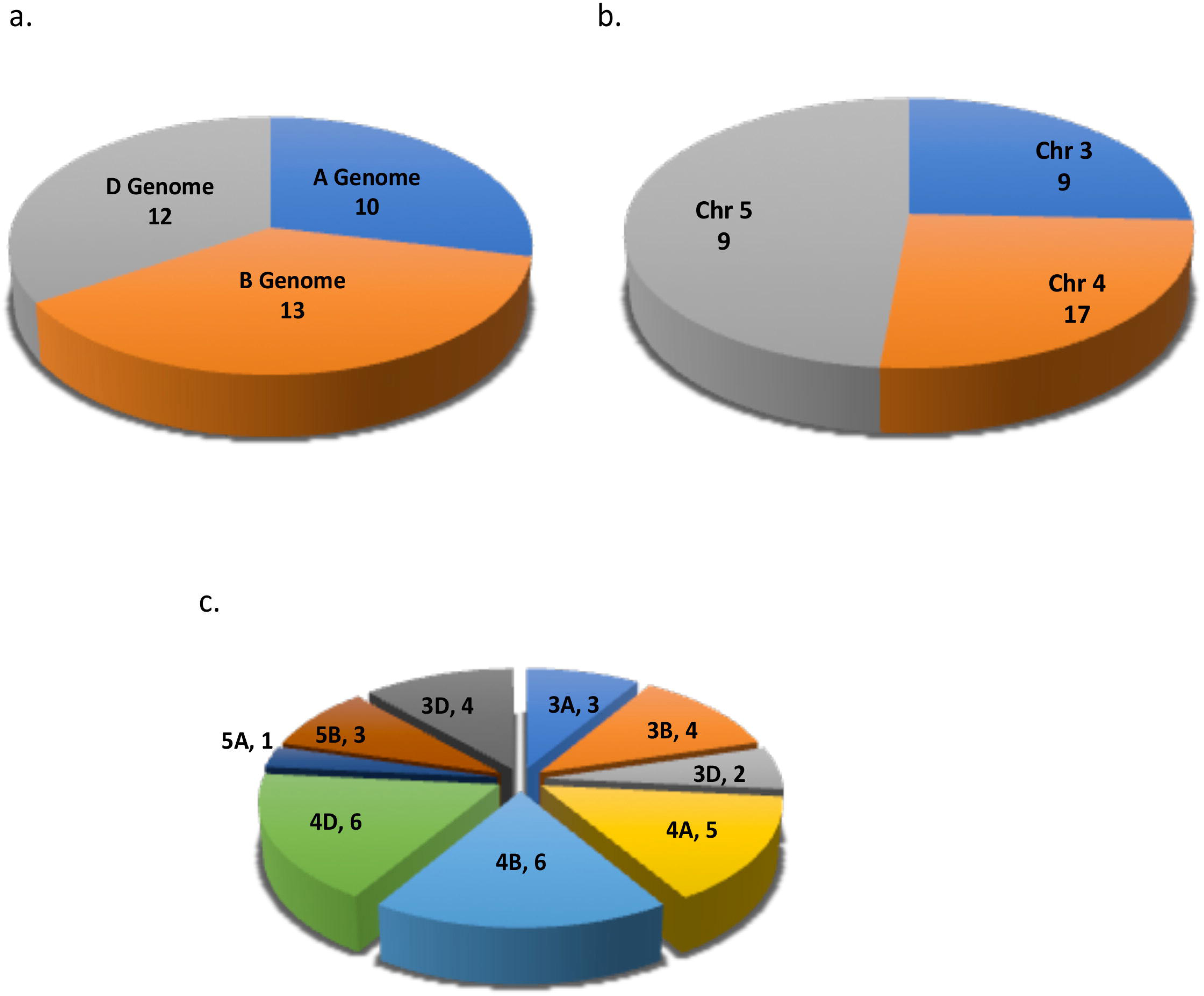
Genomic and chromosomal distribution of wheat ZIFLs on wheat genome. Distribution of thirty-five TaZIFLs across: **a**. A, B and D sub genomes. **b**. Wheat chromosomal distribution. **c**. Chromosomal distribution share on different Chromosome and genome.

**Figure 3.**
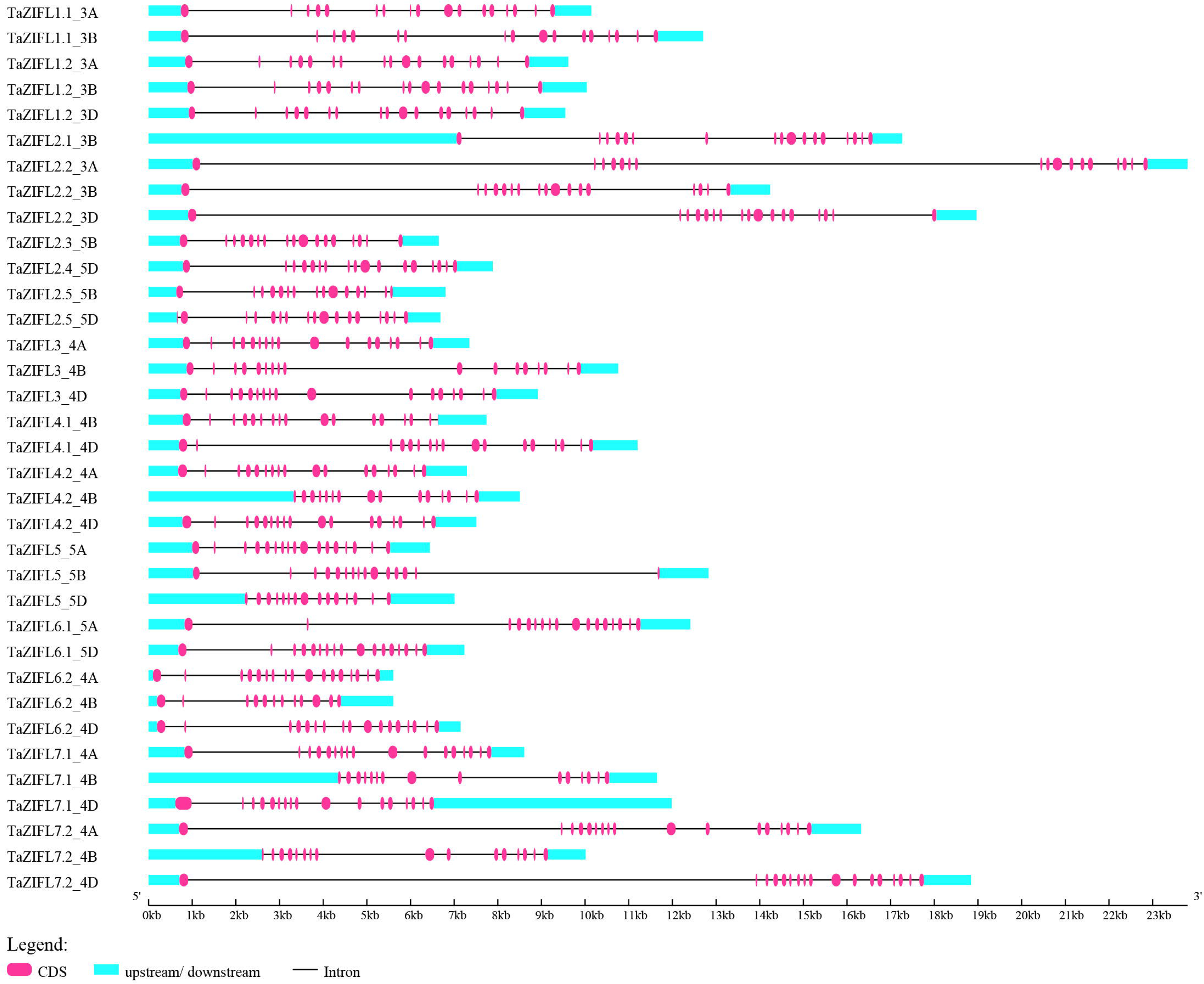
Intron exon arrangement and protein conservation. The intron-exon structure was obtained using Gene Structure Display Server (GSDS 2.0:) Yellow boxes and black lines depict the introns and exons, respectively.

### Protein motif analysis reveals presence of diverse domains

To have an understanding about the similarity, variation in motif composition and distribution of TaZIFL, 15 sequences representing each ZIFL transcript were subjected to MEME analysis. Our analysis revealed the presence of fifteen motifs (Figure 4a, Table S3). Out of fifteen motifs, six were conserved throughout all ZIFL, while some lacked few motifs. Four unique and exclusively motifs (12, 13, 14, 15) were identified, which are specific to the respective group. Motif 14 and motif 12 (Figure 4b, Table S3) are specific to TaZIFL2.1, TaZIFL2.2, TaZIFL2.3, TaZIFL2.4 and TaZIFL2.5, which indicated that TaZIFL2 members might share similar functions. Motif 14 was also present in TaZIFL6.2. Another set of unique motifs, mentioned as motif 13 and 15 was found in TaZIFL4.1 and TaZIFL4.2, which may indicate probable different function from rest of TaZIFLs. The canonical MFS signature WG[V/M/I][F/V/A/I]AD[K/R][Y/I//H/L]GRKP was present in the cytoplasmic loop between TM2 and TM3 (Figure S2, Table S4) as well S-x(8)-G-x(3)-G-P-[A/T/G]-[L/I]-G-G as anti porter signature in TM5. The results suggest that ZIFL proteins share unique signatures and high similarity indicating they are a distinct group of MFS family. Presence of conserved signatures Cysteine (Cys) -containing motif CPGC reported previously were also present in most of the wheat ZIFL proteins (Ricacahenevsky et al., 2011). The absence of these motifs was observed in TaZIFL2.5_5D, TaZIFL 3_4B, TaZIFL 4.2_4A, TaZIFL5_5D, TaZIFL7.1_4D this might be because of missing sequence information. This motif was found to be present in the cytoplasmic N-terminal loop for TaZIFL groups 2, 4, 5, 6 and in the non-cytoplasmic N-terminal loop for groups 1, 3 and 7 (Figure S2 and Table S4). Another conserved histidine (His)-containing motif PET[L/I]H showed its presence in the cytoplasmic loop between TM domains ranging from 2 and 3 to 6 and 7, with highest between 6 and 7 TM domains (Figure S2).

**Figure 4.**
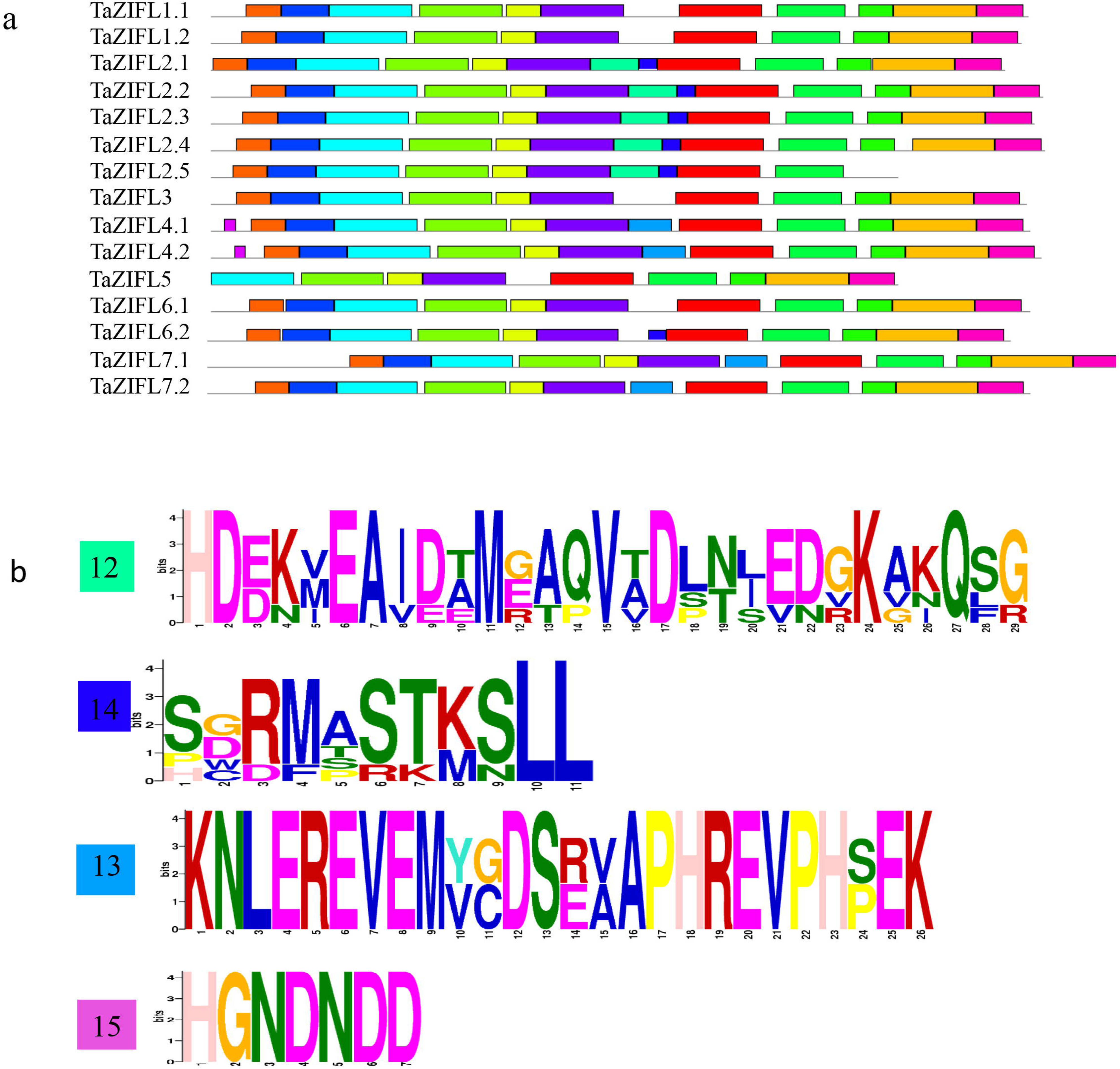
Protein sequence analysis for conserved and unique motifs in wheat ZIFL. **a.** Conserved motifs across 15 TaZIFL proteins, as obtained from MEME. Different colors represent distinct motifs. **b.** Unique motifs found through MEME for group 2, group 4, as well as TaZIFL6.2.

### Identification of conserved cis-elements in the promoters of wheat ZIFL genes

To find the molecular clues that could regulate the expression of wheat *ZIFL* transcripts, the 1.5 kB promoter region of the all identified wheat *ZIFL* genes was explored. Our analysis revealed a large number of cis-elements in the promoter of wheat ZIFL. Predominantly, the promoters were enriched with the presence of the core binding site for iron-deficiency responsive element binding factor 1 (IDEF), iron related transcription factor 2 (IRO2) and heavy metal responsive element (HMRE) (Table 5). The presence of these promoter elements suggests that wheat *ZIFL* genes might respond towards the presence of heavy metals and to important micronutrients like Fe and Zn. Interestingly, IDE1 cis-element was present on all the promoters of the respective wheat ZIFLs suggesting that they could respond to Fe limiting conditions. Few of these promoters consist of multiple such cis-elements suggesting their diverse function in plants (Table S5).

### Expression characterization of wheat ZIFL genes for their response to Zn and Fe

ZIFL are primarily known to respond towards Zn excess, therefore experiments were performed to study the gene expression of wheat ZIFL in roots and shoots. The QRT -PCR analysis suggested tissue specific expression response by wheat ZIFLs. A total of eight genes, including *TaZIFL1.2, TaZIFL2.2, TaZIFL2.3, TaZIFL4.1, TaZIFL4.2, TaZIFL5, TaZIFL6.1* and *TaZIFL6.2* showed significantly higher expression during one of the time points under Zn surplus condition (Figure 5). Of all the genes, the fold expression level for *TaZIFL4.1* was highest (∼7 fold) at 3D after treatment with respect to control roots (Figure 5a). In shoots, *TaZIFL1.1, TaZIFL1.2, TaZIFL6.1* and *TaZIFL6.2* showed significant transcript accumulation either at 3D or 6D after treatment. In our current study, a few genes like *TaZIFL 1.1, TaZIFL7.1 and TaZIFL7.2* remained unaffected by the Zn surplus condition in roots (Figure 5b). Notably, *TaZIFL1.2, TaZIFL*6.1 and *TaZIFL6.2* show enhanced transcript accumulation in both the tissues. In contrast, during our experiment the expression of a few wheat *ZIFL* genes showed down-regulated in shoots but not in roots. Our expression data under Zn surplus condition suggested the differential response by wheat *ZIFL* towards the treatment.

**Figure 5.**
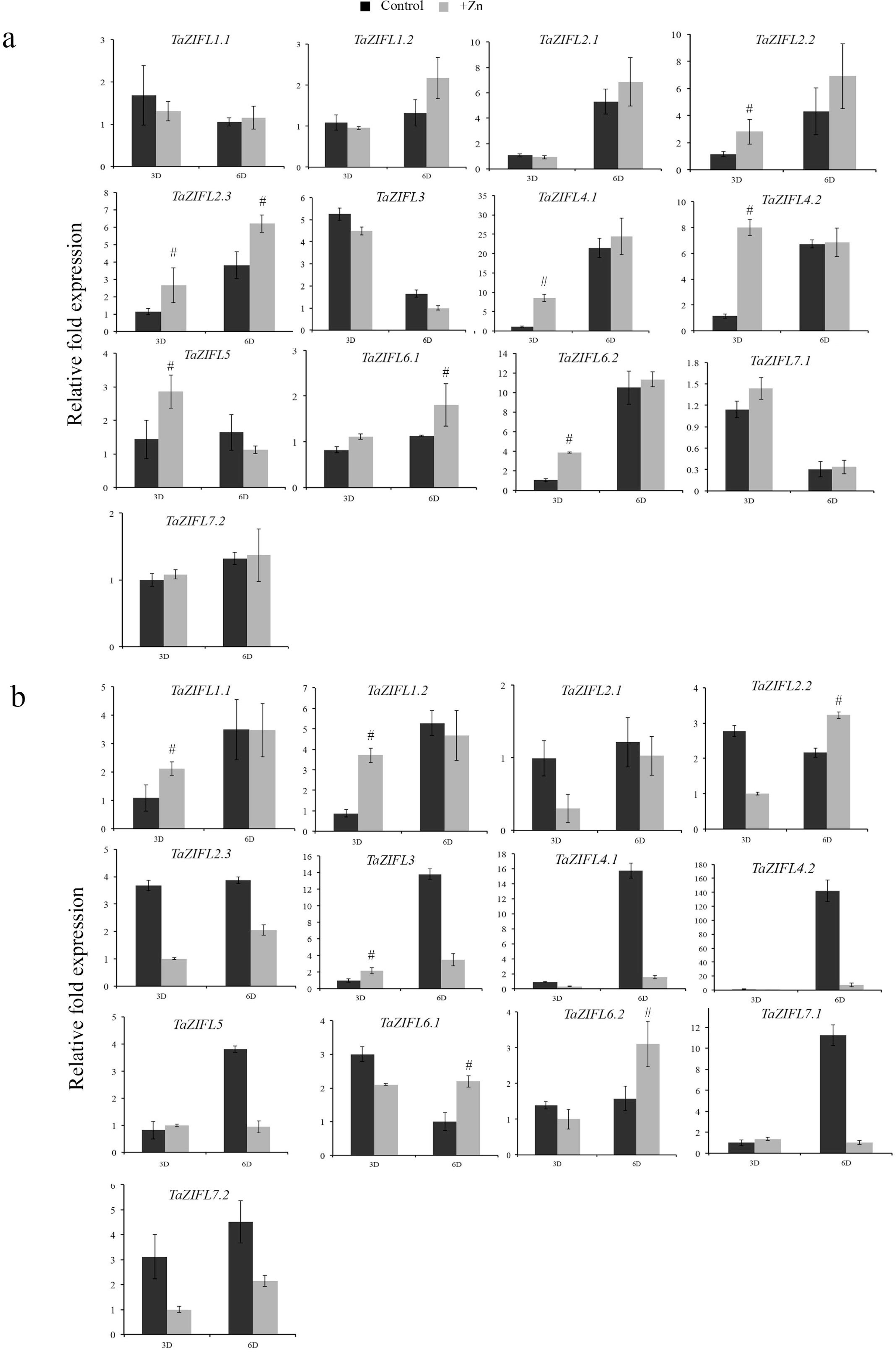
Relative gene expression levels of wheat ZIFLs under +Zn condition. Gene expression profiles of wheat ZIFL genes were studied in a) roots and b) shoots. Total RNA was extracted from the wheat seedlings subjected to three and six days of treatments +Zn. The roots and shoots samples were collected, and qRT-PCR was performed on the DNA free RNAs. A total of 2 µg of RNA was used for cDNA synthesis. C_t_ values were normalized against wheat *ARF1* as an internal control. Data represents mean of two biological replicates each treatment containing 15-18 seedlings. Vertical bars represent the standard deviation. # on the bar indicates that the mean is significantly different at p < 0.05 with respect to their respective control samples.

Previous evidences indicated that plant ZIFL genes not only respond to Zn excess, but also are also affected by the Fe limiting conditions (Haydon et al., 2012). Therefore, expression analysis of wheat *ZIFL* genes was checked in roots and shoots of seedling subjected to a Fe limiting condition. Interestingly, in the roots expression of *TaZIFL4.1, TaZIFL4.2* and *TaZIFL7.2* show up-regulation during Fe limiting condition at both at 3 and 6 days after starvation. Out of the remaining genes, *TaZIFL2.3, TaZIFL6.2* and *TaZIFL7.1* show significant transcript abundance at one-time point or the other (Figure 6a). Interestingly, in shoots *TaZIFL1.1, TaZIFL4.1, TaZIFL4.2, TaZIFL5* and *TaZIFL7.1* show up-regulation only at 3D, suggesting their coordinated response in shoots (Figure 6b). Under –Fe condition, wheat *ZIFL* genes, namely, *TaZIFL4.1* and *TaZIFL4.2* show high transcript accumulation in both roots and shoots. Remaining genes remain unaffected by the Fe stress (Figure 6b). Overall, our expression data suggested that indeed wheat ZIFL respond to the Fe limiting condition, thereby suggesting a common interlink of this gene family during Zn and Fe homeostasis.

**Figure 6.**
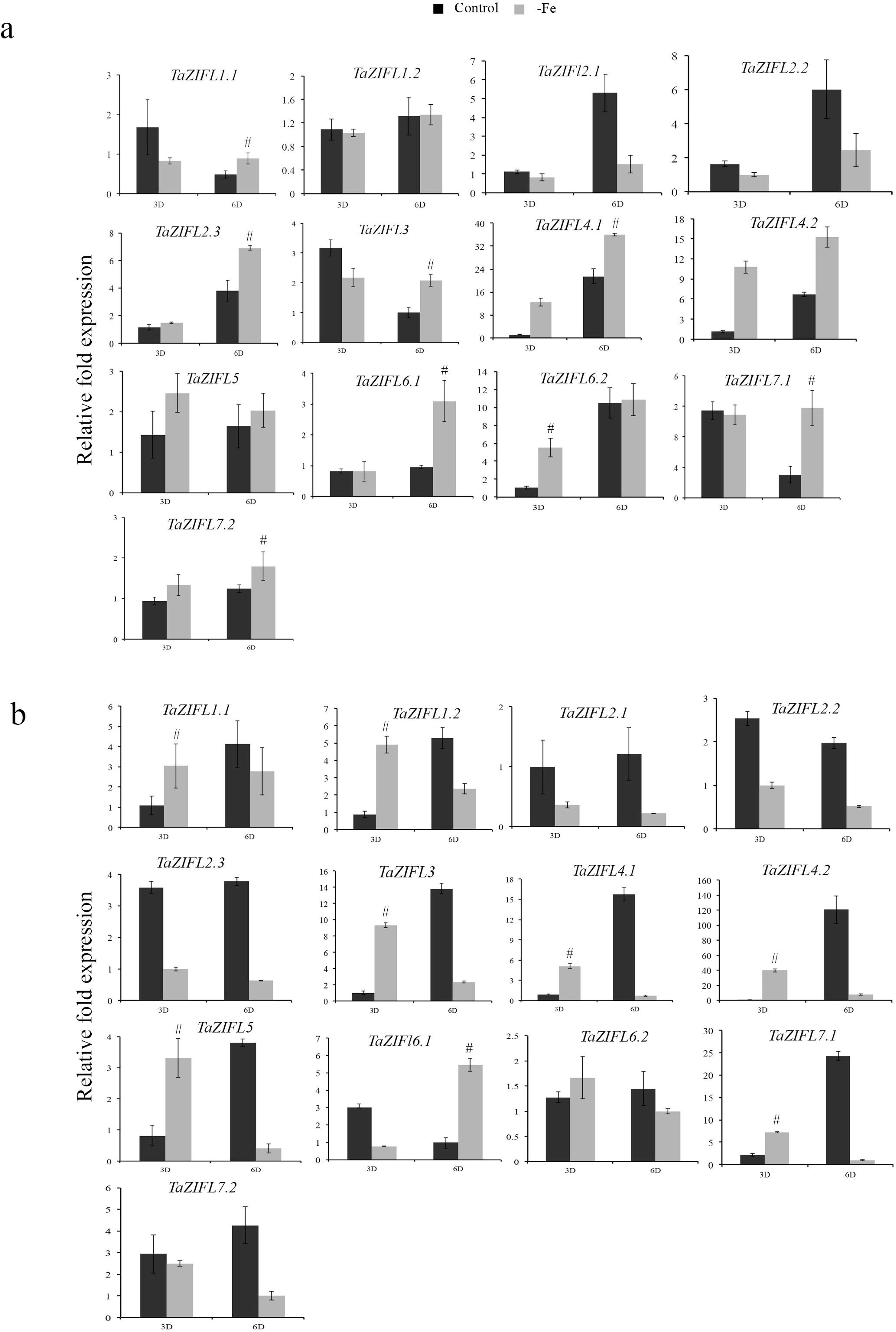
Relative gene expression levels of wheat ZIFLs under –Fe condition. Gene expression profiles of wheat ZIFL genes were studied in a) roots and b) shoots. Total RNA was extracted from the wheat seedlings subjected to three and six days of treatments. The roots and shoots samples were collected, and qRT-PCR was performed on the DNA free RNAs. A 2 µg of total RNA was used for cDNA synthesis. C_t_ values were normalized against wheat *ARF1* as an internal control. Data represents mean of two biological replicates with each treatment containing 15-18 seedlings. Vertical bars represent the standard deviation. # on the bar indicates that the mean is significantly different at p < 0.05 with respect to their respective control treatments.

### Putative wheat TOM genes showed expression response in presence of heavy metals

Our promoter analysis of wheat *ZIFL* genes indicates the presence of multiple HMRE suggesting that few of these genes could respond to the heavy metals (Supplementary Table S5). Furthermore, the phylogenetic arrangement of the wheat ZIFL proteins along with the rice suggested the corresponding candidate for the transporter of mugineic acid (TOM). Thus, based on the clade distribution for the corresponding candidate orthologs for the TOM genes for rice (TOM1-OsZIFL4, TOM2-OsZIFL5 and TOM3-OsZIFL2) are identified as TaZIFL4.1/TaZIFL4.2, TaZIFL5, TaZIFL6.1, TaZIFL6.2, TaZIFL71. and TaZIFL7.2. Due to the importance of TOM gene in micronutrient mobilization the expression of these transcripts in wheat seedlings (shoots and roots) was studied after exposure to heavy metals such as Co, Ni and Cd. During our experiment all the seedlings showed phenotypic defects when exposed to heavy metals (data not shown). Our expression analysis suggested that wheat ZIFL genes show metal specific responses. For example, *TaZIFL4.2* and *TaZIFL7.1* showed significant up-regulation in both roots and shoots when exposed to any of the metals tested (Figure 7). In contrast, the transcripts of *TaZIFL5* and *TaZIFL6.2* remained unaffected under these heavy metals. Expression of *TaZIFL7.2* showed almost no change in the presence of Ni in either yet it specifically in up-regulated in shoots when exposed to Cd or Co. Similarly, *TaZIFL6.1* showed significant upregulation only in roots upon exposure to Ni and Co (Figure 7). Overall, these data indicate the influence of specific heavy metals on the expression of wheat ZIFL genes in a tissue dependent manner.

**Figure 7.**
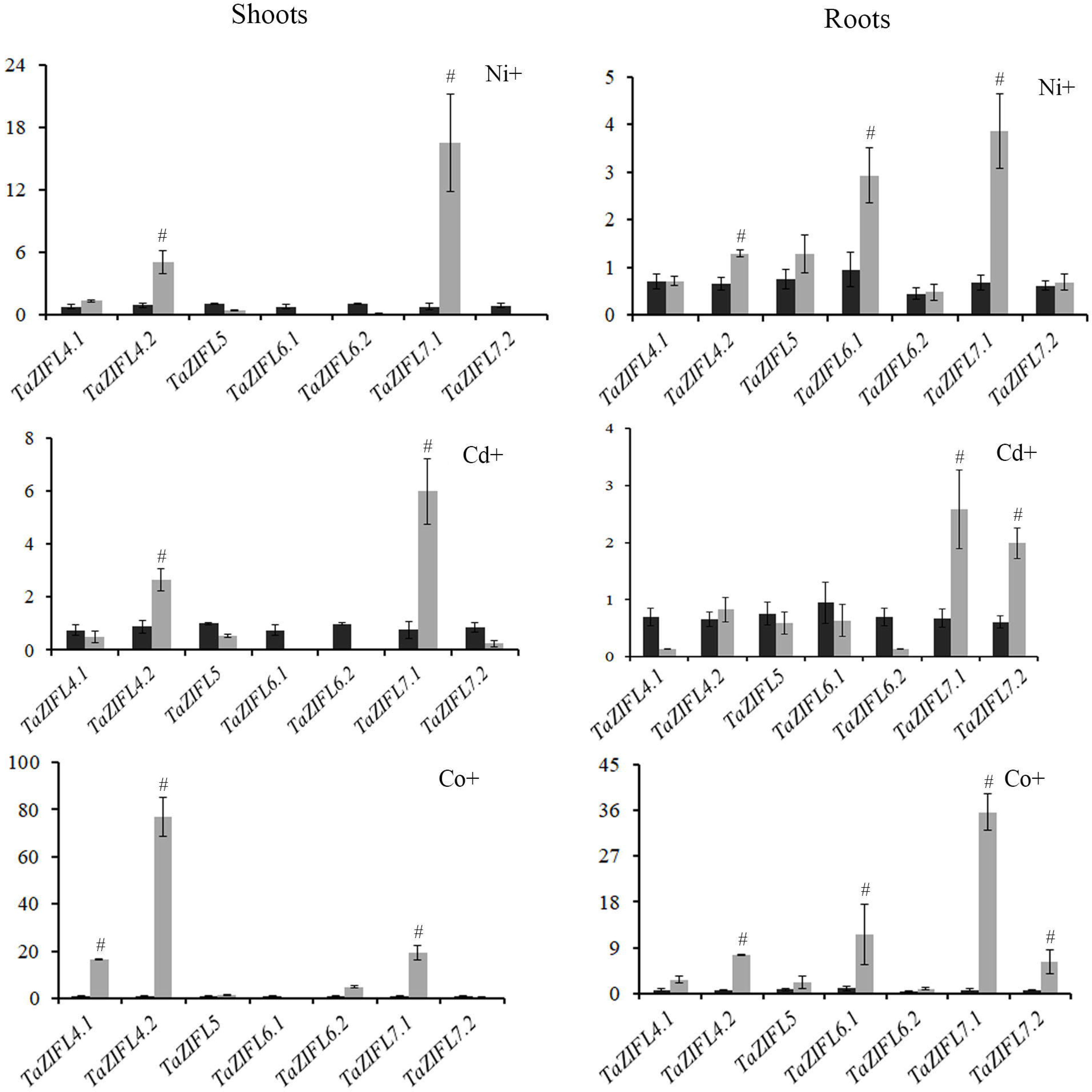
qRT-PCR expression analysis of shoots and roots of wheat seedlings exposed to multiple heavy metals (Ni, Co and cd). Five days old wheat seedlings were exposed to the mentioned heavy metals for the period of 14 days. Total RNA was extracted from the treated and control samples and 2 µg of RNA was used to construct the cDNA. C_t_ values were normalized against wheat *ARF1* as an internal control. Fold expression values were calculated relative to the control tissue of the mentioned wheat ZIFL. Vertical bars represent the standard deviation. # represent the significantly difference at p < 0.05 with respect to their respective control treatments.

### Expression of wheat ZIFL transcripts in different wheat tissues

Analysis of *ZIFL* genes was also performed in different wheat tissues and developmental stages by using transcript expression data. RNA-seq expression analysis for *TaZIFL* was also checked under different stresses. The expression values were extracted as Transcript per millions (TPM) from a wheat expression browser, expVIP (http://www.wheat-expression.com/). *TaZIFL* expression values in different tissues (aleurone-al, starchy endosperm-se, seed coat-sc, leaf, root, spike, shoot) and various developmental stages were extracted (Table S6) and were depicted as a heatmap (Figure 8). In reference to grain tissue developmental time course (GTDT) (Gillies et al., 2012), highest expression was seen for *TaZIFL1.2* (3B, 3D) and *TaZIFL5* (5A, 5B), with an increase in expression in “al” at 20 dpa and “al and se” at 30 dpa. In the expression values during grain tissue specific expression (at 12 dpa) (Pearce et al., 2014), *TaZIFL1.2* was not expressed, but like GTDT study, *TaZIFL5* (5A, 5B) was expressed in “al” as well as “se”. While for sc tissue, *TaZIFL2.2-3D* and *TaZIFL7.1* (4A, 4D) had the highest expression when compared to other *ZIFL* genes. For the tissue specific expression response *TaZIFL2.2* was abundant in spike, *TaZIFL1.2* in leaf and root. TaZIFL5 was predominantly expressed in all the tested tissue, including leaf, shoot, spike and shoot. The transcripts exclusively expressed in root were TaZIFL2.4-5D, TaZIFL2.5-5B, TaZIFL6.1-5A, and TaZIFL7.2-4D, with high induction of TaZIFL4.1 (4B, 4D), TaZIFL4.2-4A, TaZIFL6.2 (4A, 4B, 4D) for three-leaf and flag leaf stage as compared to the seedling stage. Highest expression induction was seen for TaZIFL4.2-4D. In addition, the highest expression overall in five tissues was observed for TaZIFL1.2 (3A, 3B, 3D) in leaves for seedling as well as tillering stage, TaZIFL2.2-3D in spike, TaZIFL3-4A in leaf, TaZIFL5-5A in grain, TaZIFL7.1-4D in grain and leaf.

**Figure 8.**
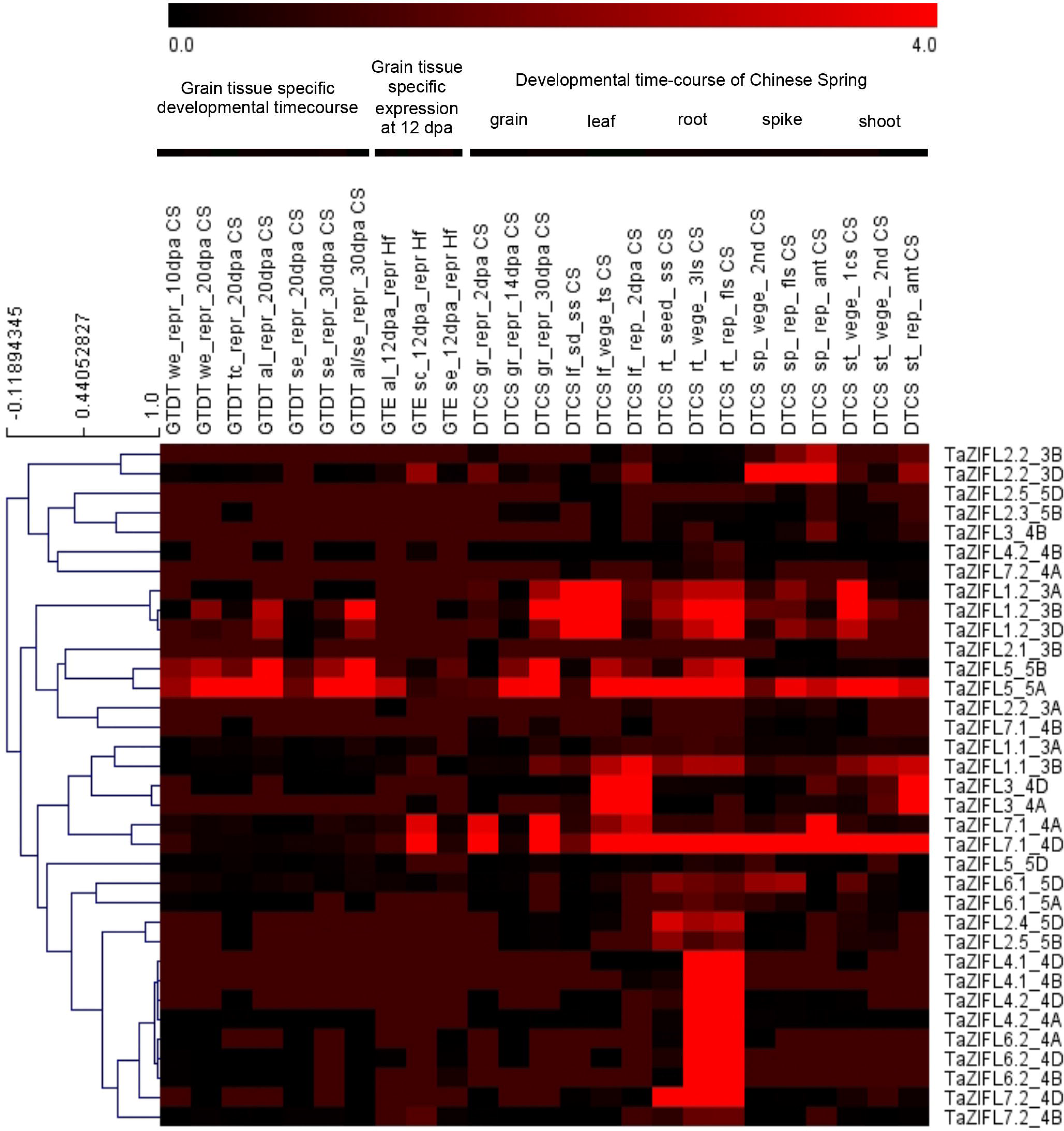
Heat map for relative expression of putative *TaZIFL* genes in different tissues and at multiple developmental stages. Heatmaps were generated using expression values from expVIP database for grain, leaf, root, spike and shoot tissues. Green to red color change depicts increase in transcript expression, as shown by the color bar.

No significant changes in the *TaZIFL* gene expression were observed for *Fusarium* head blight infected spikelets (Table S6, Figure S3). For *Septoria tritici* infected seedlings, while a ∼2-fold induction was observed for *TaZIFL1.2* (3A, 3B) after 4 days of induction, prolonged infection (13 days), resulted in its downregulation. Other ZIFLs showing changed expression were *TaZIFL1.2-3D* and *TaZIFL2.2-3A* (>2 fold up-regulation), *TaZIFL3-4B* (upto 2.7-fold downregulation). *TaZIFL1.2* (3A, 3B, 3D) were also downregulated (upto ∼4 fold) in seedlings with stripe rust infection, while only *TaZIFL1.2-3D* was downregulated under powdery mildew infection. For abiotic stress, while no major changes were observed for TaZIFLs, TaZIFL4.1 (4B, 4D) & TaZIFL4.2 (4A, 4D) were found to be significantly downregulated by ∼14-fold under phosphate starvation, while TaZIFL6.2-4D was downregulated by 3-fold (Table S6, Figure S3). Under heat, drought and heat-drought combined stress (Table S6, Figure S3), TaZIFL7.1 (4A, 4D) and TaZIFL7.2-4D were induced by upto ∼7-fold and ∼2-fold respectively, whereas TaZIFL1.2 (3A, 3B, 3D) and TaZIFL5 (5A, 5B) were downregulated by 6 and 7.5-fold, respectively. These expression data suggest that specific ZIFLs are differentially regulated under infection conditions and show perturbed expression under abiotic stresses.

## Discussion

### Wheat ZIFL proteins as putative phytopsiderophore efflux transporters

The current work was undertaken to build the inventory of wheat ZIFL. Since wheat is a hexaploid species with three genomes therefore, we expect a high number of transcripts encoding for a particular gene family. Our analysis resulted in the identification of a total of thirty-five ZIFL-like genes from hexaploid wheat. A unique observation made for the wheat ZIFLs is that all the genes are restricted to chromosome 3, 4 or 5 only. With the exception of a seven ZIFL genes, rest are localized on all the three homoeologous in the wheat genomes i.e. A, B, and D (Figure 3). This study and the previous preliminary report, led to the identification of a total of fifteen *ZIFL* with *TaZIFL1.2, TaZIFL2.2, TaZIFL3, TaZIFL4.2, TaZIFL5, TaZIFl6.2, TaZIFL7.1* and *TaZIFL7.2* showing the presence of all the homoeologous (homoalleles) genes (Pearce et al., 2014). All the wheat ZIFLs belongs to the MFS superfamily, thereby containing the canonical ZIFL MFS signature and MFS antiporter sequence (Figure S3). Given the high sequence homology, they are named from TaZIFL1 to TaZIFL7 according to their clad distribution in the phylogenetic tree that corresponds to the rice genes. Previously, in rice multiple TOM genes were identified as protein belonging to the ZIFL sub-family. Subsequent characterization of these rice ZIFL genes led to the identification of functionally active *OsTOM1, OsTOM2* and *OsTOM3* (Nozoye et al., 2011; Nozoye et al., 2015). Therefore, based on the phylogenetic arrangement and previous characterization in rice, the corresponding homolog for the putative functional wheat TOM could be *TaZIFL4.1/4.2, TaZIFL5* and *TaZIFL7.1/7.2*. Based on our analysis, wheat ZIFL proteins show localization on the PM, except for TaZIFL4.2-B and TaZIFL5-D. These two wheat ZIFL predicted proteins are putatively localized on vacuolar membrane, thereby making them a strong candidate in a quest to identify novel membrane transporters for micronutrient storage in the cell organelles. In general, very small numbers of *ZIFL* genes are being reported from dicot species like *Arabidopsis* and *Vitis vinifera* and *Populus trichocarpa* (Ricachenevsky et al.,2011). Given the complexity of the wheat genome one could certainly anticipate the presence of multiple possible putative phytopsiderophore efflux transporters that needs to be functionally characterized in the near future. In rice, the high ZIFL numbers in rice could be accounted due to the lineage-specific expansion of the gene family (Ricachenevsky et al., 2011). Overall, our analysis identified highest number of *ZIFL* genes reported till date from any monocot species.

### Wheat ZIFL genes display overlapping gene expression

Plants undergoing metal stress result in series of signaling events that largely includes reprogramming of transcripts that could help them to overcome the toxicity. In plants, excess of Zn also results in the generation of reactive oxygen and nitrogen species (Feigl et al., 2015). The abundance of multiple membrane proteins is increased by the presence of either excess Zn or Fe deficiency. Likewise, in the previous studies, our data also confirmed the overlapping expression response of wheat ZIFL genes (Briat et al., 2015). Additionally, under Fe limiting conditions, induction of Zn responsive genes could be an important step towards limiting the non-specific transport activity of transporters which are primarily induced for Fe deficiency. ZIFL are well known for their response towards the presence of excess Zn. During our study, we also observed multiple wheat ZIFLs that responded to excess Zn, either in shoots or roots in a temporal manner. Interestingly, most of the wheat ZIFL genes show the presence of iron responsive cis-element IDE1. IDE1 is one of the primary cis-elements that respond to Fe limiting conditions (Kobayashi et al., 2007). Therefore, we studied the expression of wheat ZIFLs under –Fe condition. Multiple ZIFLs showed specific response towards Fe deficiency, suggesting that iron deficiency response by ZIFLs could be mediated by certain transcription factors like IDE1 that are highly specific for Fe homeostasis. Expression of the few of the ZIFLs like *TaZIFL1.*1, *TaZIFL1.2, TaZIFL4.2* and *TaZIFL6.2* were also affected by both Fe and Zn. These results suggest that few of these ZIFLs might be involved in the overlapping pathways of Fe and Zn homeostasis. Such partial overlaps of Zn and Fe homeostasis was reported earlier and is also evident from our work. This may also lead to a speculation for the sharing of a common network of transcription factors related to Fe and Zn interactions. In our study, the promoters of *TaZIFL1.2* and *TaZIFL2.3* showed the presence of IRO2 binding domains that has been previously speculated to be the link between Fe and Zn homeostasis (Ricachenevsky et al., 2011). Nonetheless, only *TaZIFL1.2* respond to –Fe and +Zn stress that is restricted only to shoots. Previously, it was also shown that expression of Arabidopsis ZIF1 remained unaffected in the presence of sub-inhibitory levels of Cd or Cu (Haydon and Cobbette, 2007). Based on the expression response of ZIFL genes during in the presence of heavy metals it seems that *TaZIFL5* and *TaZIFL6.2* could be one of the best candidate genes for the further studies, as both the genes remained unaffected. Nevertheless, careful selection of candidate gene must be done to minimize the cotransport of other undesired metals during micronutrient uptake.

In addition to their anticipated role in Fe and Zn homeostasis, their role in root development has been also proven. Plant ZIFL transporters have been reported to regulate stomatal movements by means of polar auxin transport, thereby modulating potassium and proton fluxes in *Arabidopsis* (Remy et al., 2013). Analysis of the expVIP data suggested high expression of wheat ZIFLs (TaZIFL7.1-4A and 4D) under drought condition. Earlier, *zifl-1* and *zifl-2* mutants of *Arabidopsis* showed hypersensitivity to drought stress by disruption of guard cells activity (Remy et al., 2015). *TaZIFL1.2* is the wheat transporter showing highest expression in leaf and highest homology with *AtZIFL1*, thereby belonging to the same clade in the phylogeny tree. In contrast to the expected function of ZIFL in Fe and Zn homeostasis, a putative role has been demonstrated in plant defense. Maize ZIFL referred as Zm-mfs1 was high induced during plant defense and has been implicated its role for export of antimicrobial compounds during its interaction with the bacterial pathogen (Simmons et al., 2003). In our study, a strong expression of multiple *ZIFL* genes was observed when infected with multiple pathogens suggesting its important role in providing resistance against fungal pathogens (Figure S3).

This study concludes that ZIFL transporters are the important players during the crosstalk of Fe and Zn homeostasis. With the recent evidences regarding its role as a potential transporter for nicotinamine and mugineic acids in roots, ZIFL are certainly a priority candidate for uptake and remobilization of micronutrients in the cereal grains like wheat.

## Conclusion

This is the first comprehensive study that resulted in identification of fifteen putative ZIFL genes from hexaploidy wheat at the homoeolog level. These are the highest number of ZIFL genes reported in plant system till date. Wheat ZIFL were characterized for their expression response in seedlings exposed to excess Zn and Fe starvation. The contrasting expression of these ZIFL in presence of heavy metals suggested their functional redundancy and pinpoint importance of a few for further functional validation. Overall, we identified few candidate ZIFL from hexaploid wheat that could be the important target to address new means to enhance micronutrient uptake.

## Abbreviations used

CCC1: Ca^2+^-sensitive cross complementer 1
YSL: yellow-stripe like protein
ZIFL: Zinc-induced facilitator-1 like gene
CDD: conserved domain database
MFS: Major facilitator superfamily
TM: trans-membrane
HMRE: heavy metal responsive element
IRO2: iron related transcription factor 2
IDEF: iron-deficiency responsive element binding factor 1
QRT-PCR: quantitative real time polymerase chain reaction
TOM: transporter of mugineic acid
TPM: transcript per millions
PM: plasma-membrane

## Methods

### Identification of the MFS-1 family in wheat

To identify the potential members of *ZIFLs* from MFS_1 transporter family in wheat genome, we used two independent approaches. In the first approach, the Pfam number (PF07690) for MFS_1 was used and sequences were extracted from wheat using the Ensembl wheat database. As a complementary approach, known sequences of *ZIaFL* genes from *A. thaliana, O. sativa* and *B. distachyon* were retrieved and used for BLAST analysis against the wheat databases: Ensemble (http://plants.ensemble.org/Triticum_aestivum/) to retrieve the sequences. The identified *MFS_1* superfamily was validated through the domain search in CDD-NCBI database (https://www.ncbi.nlm.nih.gov/Structure/cdd/wrpsb.cgi) [33].

### Identification, classification, and chromosomal distribution of wheat *ZIFL* genes

To identify putative *TaZIFLs* genes among MFS_1 superfamily sequence, phylogenetic tree was constructed with known ZIFLs from different plants that separated the ZIFL cluster from the rest of the MFS_1 superfamily members. This distribution of genes in ZIFL cluster was validated through the presence of ZIFL specific signature sequences. To construct the phylogenetic tree one hundred seventy-nine wheat MFS_1 superfamily protein sequence was retrieved from Pfam no. (PF07690). In addition to that, thirteen-protein sequence of ZIFL from *O. sativa*, five from *Zea mays* and three from *Arabidopsis* were used. Identified TaZIFL genes were named according to their corresponding rice orthologs having maximum number of ZIFL reported. The name indicates corresponding ortholog from rice followed by chromosomal and genomic location e.g. *TaZIFL3-4A, TaZIFL3-4B* and *TaZIFL3-4D* represent three homoeologous of *TaZIFL* gene in chromosome four of all the three A, B and D sub-genomes and it is an ortholog of *OsZIFL3.* Phylogenetic analysis was also done with identified ZIFL proteins of wheat and other plant species (*O. sativa, Z. mays* and *Arabidopsis)* to know the evolutionary relationship among them. All the proteins were aligned through MUSCLE algorithm and a rooted phylogenetic tree was used to construct with the Neighbor-joining (NJ) method using MEGA7 software with 1000 bootstrap replicates [34]. To determine the distribution of *ZIFL* genes in the wheat chromosomes, the position for each *ZIFL* genes was obtained using wheat Ensembl database (ftp://ftp.ensemblgenomes.org/pub/plants/release-34/fasta/triticum_aestivum).

### Analysis of conserved domains, gene arrangements and subcellular localization of wheat ZIFL

The divergence and conservation of motifs in wheat ZIFL proteins were also identified by using MEME (Multiple Expectation Maximization for Motif Elicitation) program version 5.0.2 (http://meme-suite.org) [35] with maximum motif width, 50; maximum number of motifs,15; and minimum motifs width. Gene Structure Display Server (GSDS 2.0) was used to analyze the gene structure. Individual wheat ZIFL’s CDS and corresponding genomic DNA were aligned to identify the intron-exon arrangement. Using Expasy Compute PI/MW online tool (http://us.expasy.org/tools/protparam.html) the predicted isoelectric points and molecular weights of putative TaZIFLs were calculated. To predict terminal ends and number of transmembrane domains, TMHMM (http://www.cbs.dtu.dk/services/TMHMM/) was utilized. The putative protein sequences of ZIFL genes were further *in silico* analyzed predict their subcellular localization by WoLF PSORT (https://wolfpsort.hgc.jp/) prediction program [36].

### Plant material and Fe, Zn and heavy metals treatments

For giving various zinc and iron treatment (+Zn) and (-Fe), seeds of *Triticum aestivum* cv. C306 was used. The seeds were washed with double autoclaved water for the removal of dirt followed by surface sterilization with 1.2% Sodium hypochlorite prepared in 10% ethanol. Seeds were stratified by keeping them overnight at 4 °C on moist Whatman filter papers in a Petri dish. The stratified seeds were further allowed to germinate at room temperature. Healthy seedlings were transferred to phytaboxes (12-15 seedlings/phytabox) and grown in autoclaved water in growth chamber for 5 days. Five days old plantlets were subjected to different Zn and Fe conditions and grown in Hoagland media [37] supplemented with either 2.5 µM of Fe (III) EDTA for iron deficient condition (-Fe) or 200µM of ZnSO_4_.7H_2_O for zinc surplus experiment (+Zn). Seedlings grown in the Hoagland media containing 20 µM of Fe (III) EDTA and 2 µM of ZnSO_4_.7H_2_O were used for control experiments. The plantlets (5 days post germination) tissues were collected after 3 and 6 days (D) post treatments along with the respective controls. Every alternate day, the seedlings in the phyta-boxes were supplemented with the fresh media. For heavy metal treatment the 5 days old plantlets were subjected to cadmium (50 µM CdCl_2_), Cobalt (50 µM CoCl_2_), Nickel (50 µM NiCl_2_) treatment. The tissue sample (root and shoot) were collected after 15 days of treatment. (These experiments were repeated twice, frozen in liquid nitrogen and store at −80°C.

### RNA isolation, cDNA preparation and qRT-PCR analysis

Total RNA was extracted from harvested roots and shoot samples using TRIZOL RNA extraction method. Turbo DNAfree kit (Invitrogen) was used to remove the genomic DNA. RNA samples were then quantified on nanodrop and subsequently, 2 µg of total RNA was used to prepare cDNA by using SuperScript III First-Strand Synthesis System (Invitrogen). For expression analysis qRT-PCR primers were designed from the conserved region of respective *TaZIFL* genes. (Table S7). Amplicons arising from these primers were also processed for sequencing to avoid any cross amplifications of the *ZIFL* genes. 10X diluted cDNA and SYBER Green I (QuantiFast^®^ SYBR^®^ Green PCR Kit, Qiagen) was used to perform qRT-PCR on 7500 Fast Real-Time PCR System (Applied Biosystems, USA). The relative mRNA abundance was normalized with wheat *ARF* (*ADP-Ribosylation Factor*, AB050957.1; [38, 39]. The relative expression was calculated through delta-delta CT-method (2^-ΔΔCT^) [40]. The statistical significance of expression data was determined using student’s t-test (p-value <0.05).

### *In-silico* expression analysis

For *In-silico* expression analysis 35 *TaZIFL* genes were selected and wheat expression browser, expVIP [http://www.wheat-expression.com/] was used to extract the expression values in the form of TPMs. These values were then used to build heatmaps using MeV software [http://mev.tm4.org/]. While absolute values were used for development and tissue-specific data, fold change values were used for stress conditions where a gene was taken to be upregulated if fold change was greater than 2 and downregulated if less than 0.66. For abiotic stress, expression for two studies, phosphate starvation [41] and heat, drought and heat-drought stress [42] was studied. In case of biotic stress, the studies considered were [43–45].

## Supporting information

Supplementary Images S1_S3

## Declarations

### Ethics approval and consent to participate

Not applicable

### Consent to publish

Not applicable

## Availability of data and materials

Not applicable

## Funding

This research was funded by the NABI-CORE grant to AKP

### Acknowledgments

The authors thank Executive Director, NABI for facilities and support. This research was funded by the NABI-CORE grant to AKP. Thanks to the International Wheat Genome Sequencing Consortium for providing the high-quality wheat genome resources.

## Author contribution

AKP and SS conceptualized and planned the work. SS, AK, JK and AKP designed the study. SS and AK performed all the experiments. GK and SS performed the Bioinformatics work. SS, AK and AKP analyzed the data. AKP, GK, JK and SS wrote the manuscript. All the authors reviewed and edited the final version of the manuscript.

## Legends for the Supplementary Figures

**Figure S1** Neighbor-Joining (NJ) tree for MFS_1 family of proteins from *Oryza sativa, Brachypodium distachyon, Zea mays* and *Triticum aestivum* constructed using MEGA7.0 software with a bootstrap replicate value of 1000. The phylogenetic tree shows ZIFL proteins clustering into a distinct clade within the MFS family.

**Figure S2** Showing MEGA alignments for signature sequences specific for ZIFLs, namely, ZIFL MFS signature motif, the anti-porter signature, Cys-containing, His-containing signatures.

**Figure S3** Relative expression of putative *TaZIFL* genes under various abiotic and biotic stresses: (A) Heat maps of *TaZIFL* genes generated using fold change values obtained after processing of TPM values from wheat expression database expVIP under various abiotic (Phosphate starvation, Drought, Heat and Drought-Heat) (B) Biotic stresses (*Fusarium* heat blight, *Septoria tritici*, stripe Rust and Powdery mildew). Color bar shows the fold change values, thereby green color represents downregulation, black represents no change and red color represents upregulation.

## Legend for Supplementary Tables

**Table S1:** List of 179 wheat sequence IDs extracted for MFS_1 family, Pfam ID: PF07690 using ensembl wheat database

**Table S2:** Detailed information for 35 putative wheat ZIFLs identified. Table includes gene IDs, chromosomal locations, CDS and protein lengths, molecular weight and pI for each of the obtained putative *TaZIFLs*.

**Table S3:** Conserved motifs identified in putative TaZIFLs using MEME. The consensus sequence logo, e-values and the number of sequences in which each motif was found are listed.

**Table S4:** ZIFL Signature motifs (Cys, His, MFS and MFS antiporter) and their locations in all TaZIFLs. The motif positions were mapped to the TM-HMM predicted TaZIFL protein structures to obtain the location of motifs.

**Table S5:** Cis-elements of wheat ZIFLs along with their positions in the promoter. HMRE-heavy metal responsive element, IRO2-iron related transcription factor 2, MRE: metal responsive element, IDE1: iron-deficiency responsive element binding factor 1.

**Table S6:** Table listing the details and expression values extracted from expVIP for different developmental stages and tissues, as well as fold change values obtained after processing the TPM values for the stress conditions (Abiotic stress: phosphate starvation; heat, drought, combined heat-drought stress, & Biotic stress: Fusarium heat blight, *Septoria tritici*, stripe Rust and Powdery mildew).

**Table S7:** List of the primers used during the current study.

