## Supplementary Images S1_S3 for "Overlapping transcriptional expression response of wheat zinc-induced facilitator-like transporters emphasize important role during Fe and Zn stress"

### Slide 1
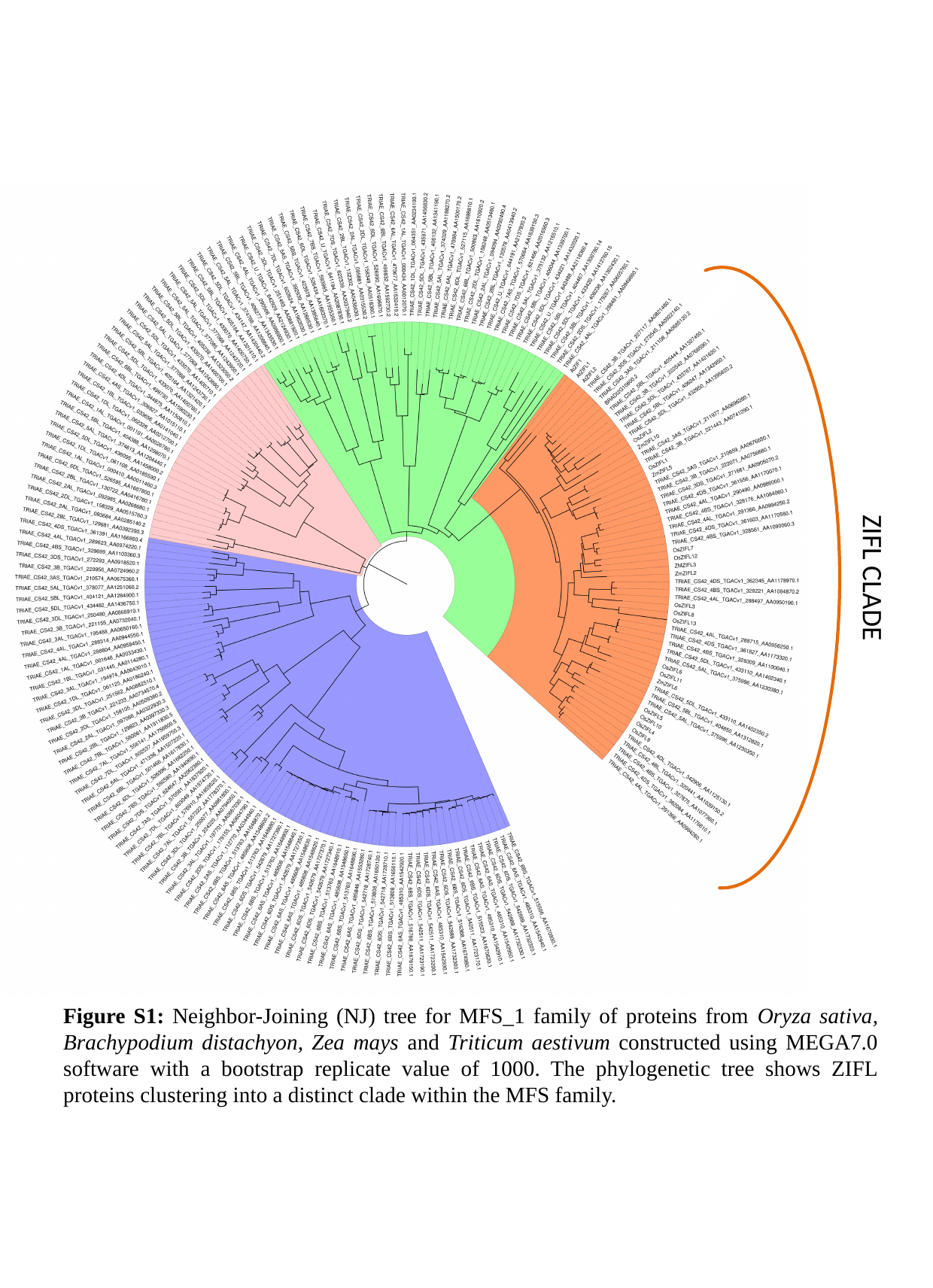

ZIFL CLADE
Figure S1: Neighbor-Joining (NJ) tree for MFS_1 family of proteins from Oryza sativa, Brachypodium distachyon, Zea mays and Triticum aestivum constructed using MEGA7.0 software with a bootstrap replicate value of 1000. The phylogenetic tree shows ZIFL proteins clustering into a distinct clade within the MFS family.

### Slide 2
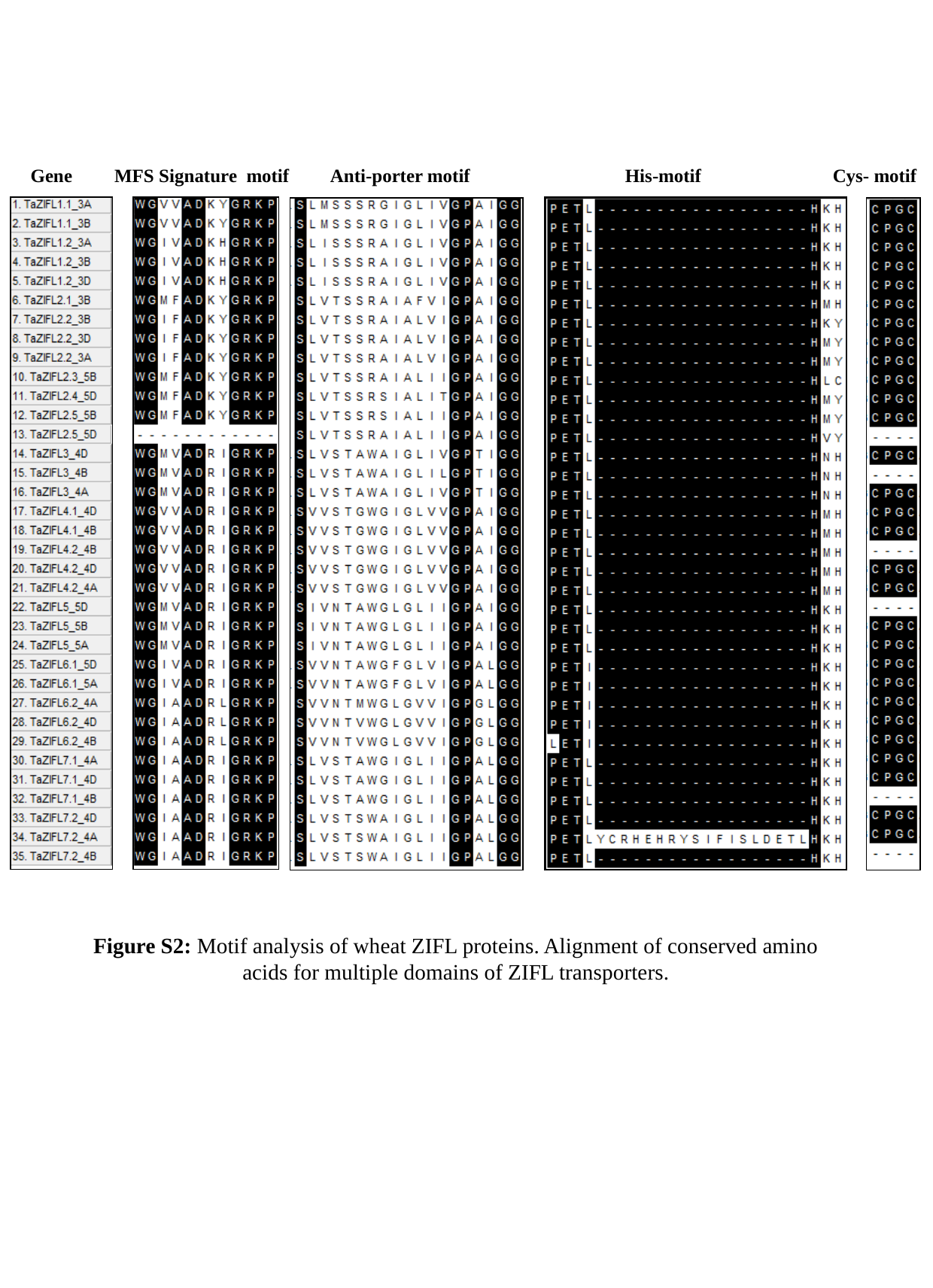

Gene MFS Signature motif Anti-porter motif His-motif Cys- motif
Figure S2: Motif analysis of wheat ZIFL proteins. Alignment of conserved amino acids for multiple domains of ZIFL transporters.

### Slide 3
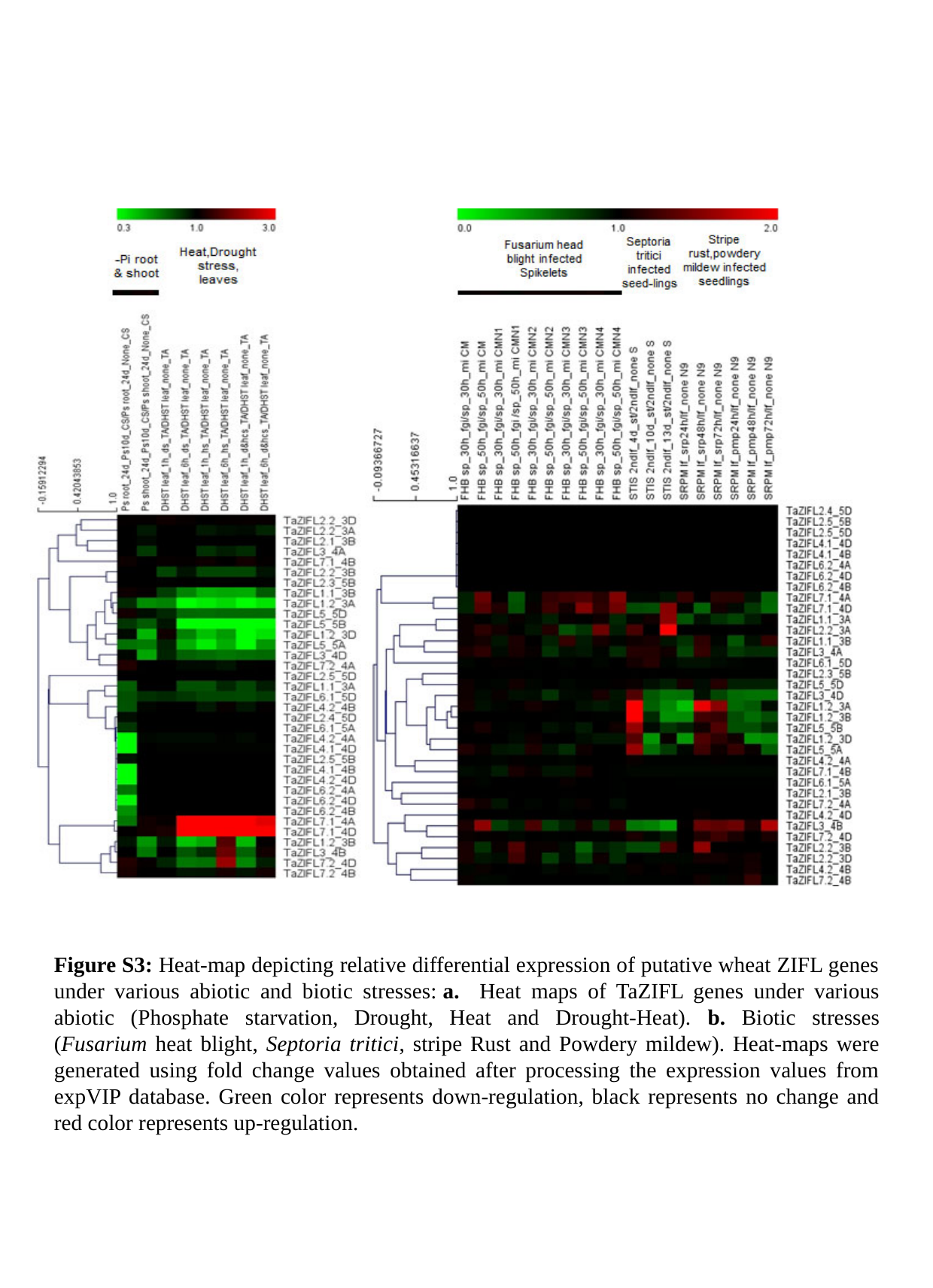

Figure S3: Heat-map depicting relative differential expression of putative wheat ZIFL genes under various abiotic and biotic stresses: a. Heat maps of TaZIFL genes under various abiotic (Phosphate starvation, Drought, Heat and Drought-Heat). b. Biotic stresses (Fusarium heat blight, Septoria tritici, stripe Rust and Powdery mildew). Heat-maps were generated using fold change values obtained after processing the expression values from expVIP database. Green color represents down-regulation, black represents no change and red color represents up-regulation.
