## Supplemental Table S3 for "Overlapping transcriptional expression response of wheat zinc-induced facilitator-like transporters emphasize important role during Fe and Zn stress"

**Supplementary Table S3:** Conserved motifs identified in TaZIFL proteins by MEME search. Table shows the consensus sequence logo, E-values and the number of proteins in which each motif is found.

| **Motif No.** | **Motif** | **E-value** | **Sites out of 14** |
| --- | --- | --- | --- |
| 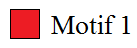 | 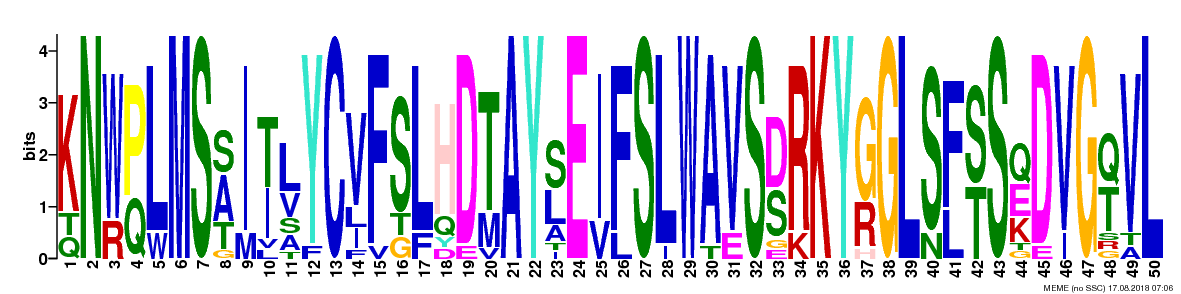 | 2.7e-453 | 14 |
| 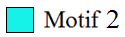 | 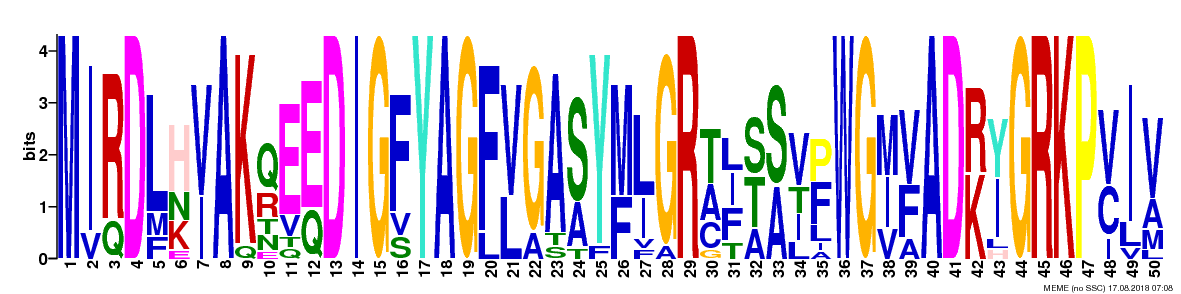 | 2.6e-430 | 14 |
| 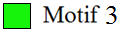 | 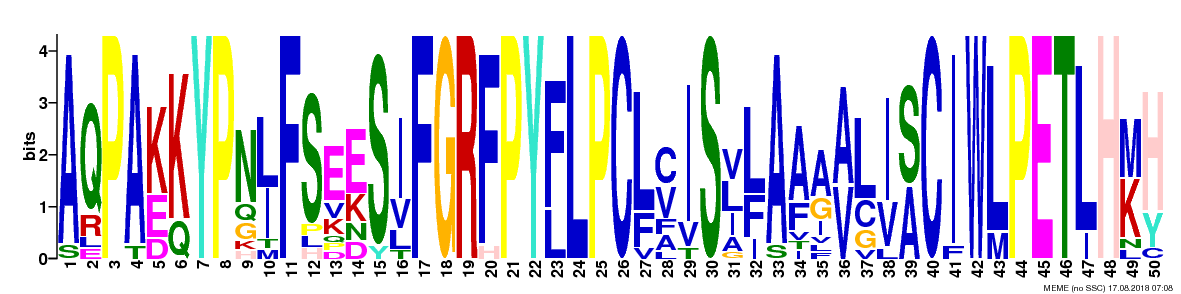 | 3.5e-430 | 14 |
| 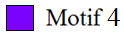 | 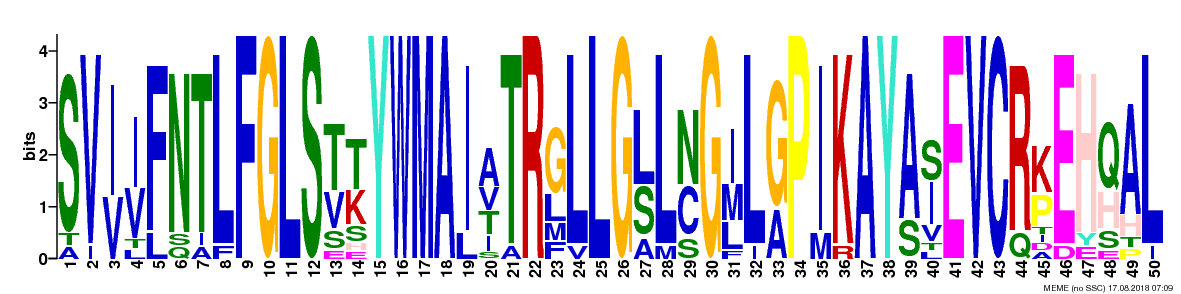 | 7.5e-426 | 14 |
| 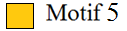 | 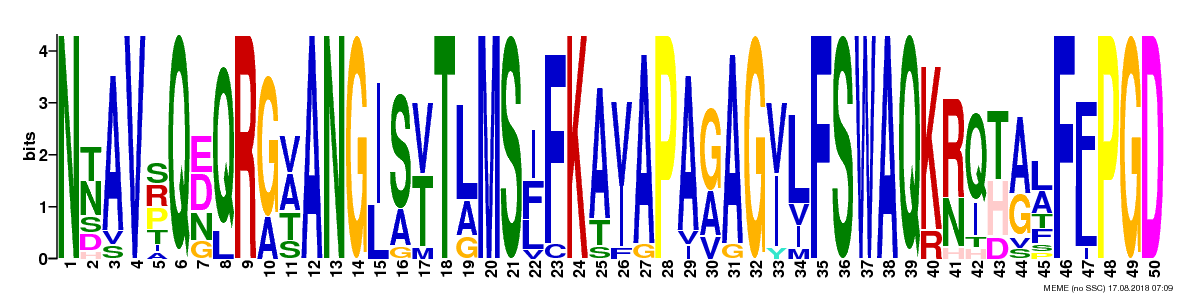 | 1.2e-385 | 13 |
| 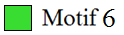 | 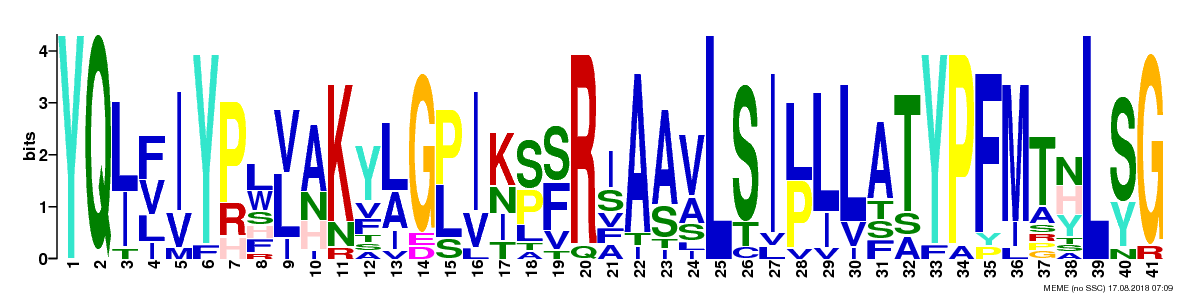 | 3.6e-217 | 14 |
| 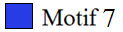 | 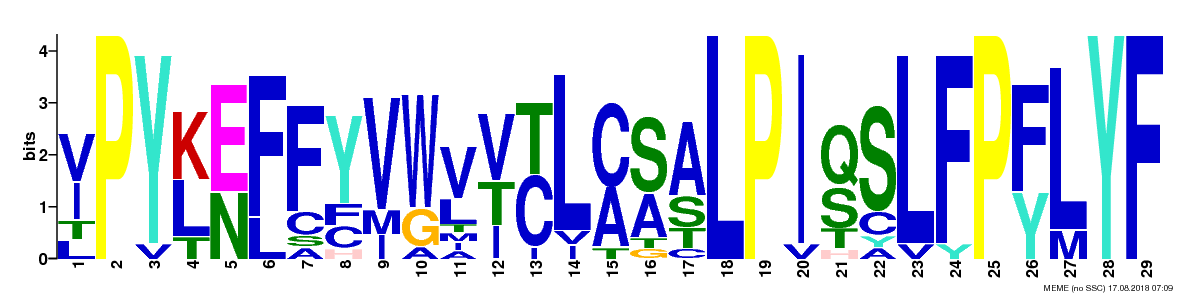 | 8.6e-171 | 13 |
| 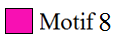 | 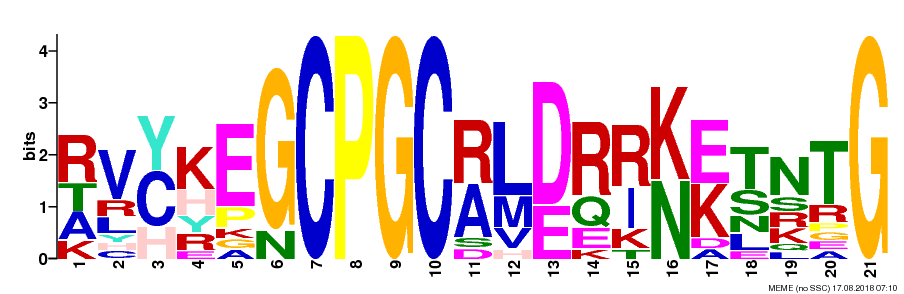 | 6.3e-121 | 13 |
| 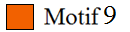 | 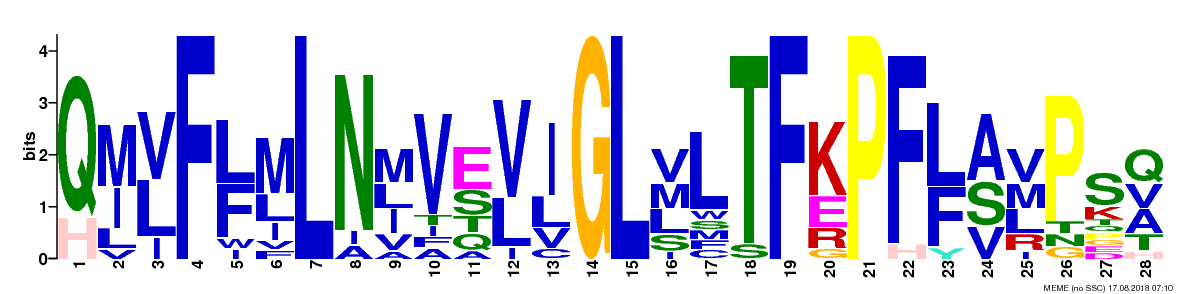 | 9.4e-116 | 13 |
| 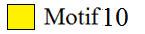 | 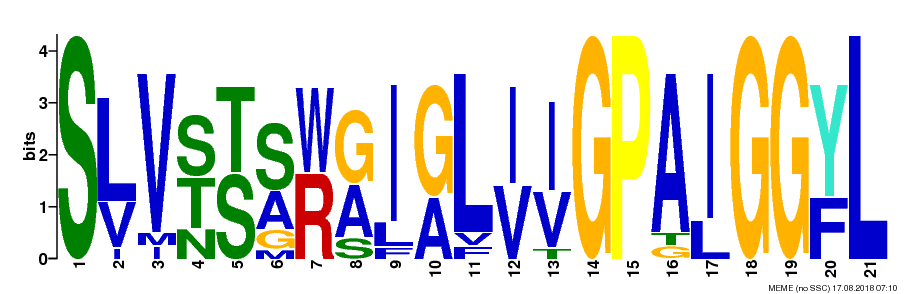 | 1.2e-115 | 14 |
| 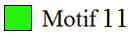 | 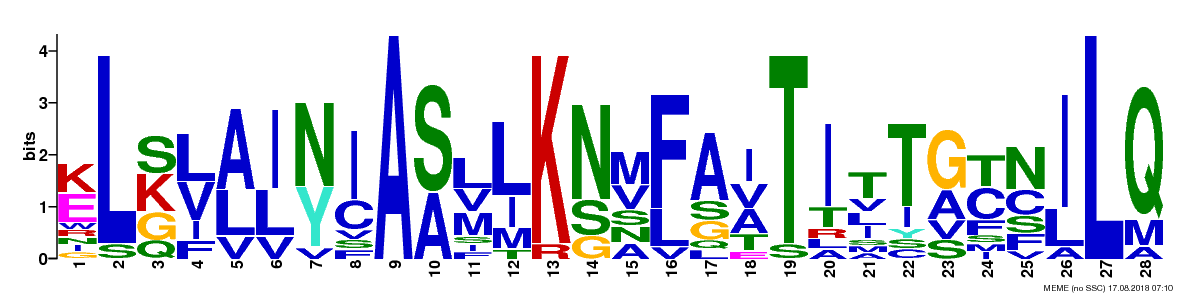 | 3.7e-078 | 13 |
| 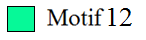 | 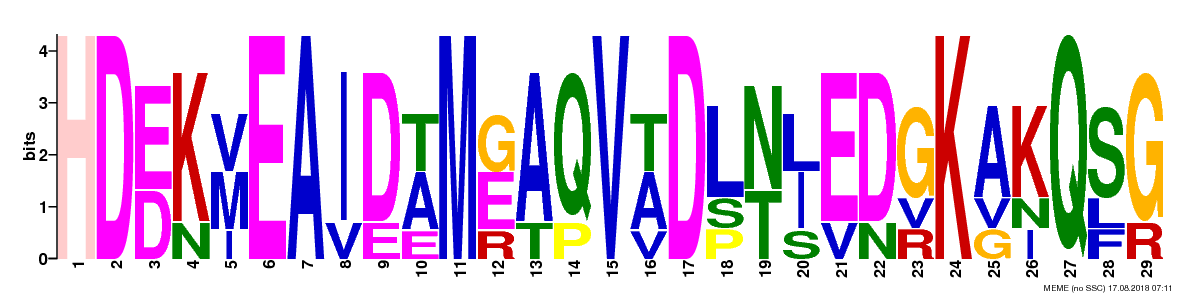 | 1.9e-051 | 5 |
| 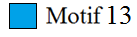 | 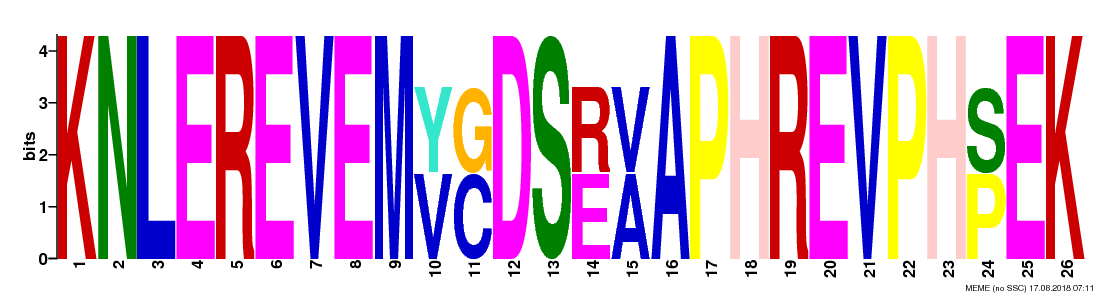 | 1.4e-010 | 2 |
| 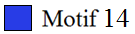 | 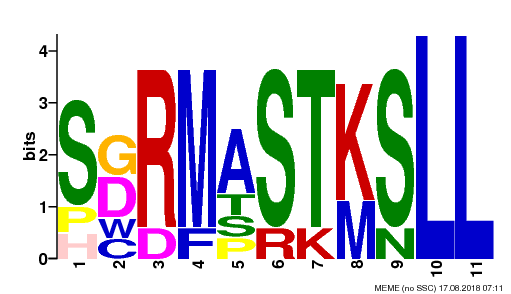 | 4.6e-006 | 6 |
| 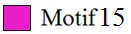 | 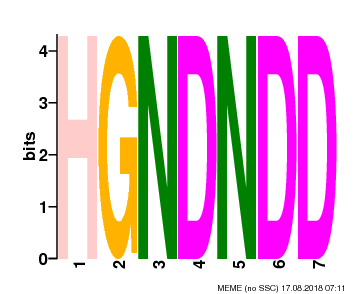 | 1.5e-003 | 2 |
